# PaxtoolsR: Pathway Analysis in R Using Pathway Commons

**DOI:** 10.1101/021105

**Authors:** Augustin Luna, Özgün Babur, Bülent Arman Aksoy, Emek Demir, Chris Sander

**Affiliations:** Computational Biology Center, Memorial Sloan Kettering Cancer Center, New York, NY 10065, USA

## Abstract

**Purpose:** PaxtoolsR package enables access to pathway data represented in the BioPAX format and made available through the Pathway Commons webservice for users of the R language. Features include the extraction, merging, and validation of pathway data represented in the BioPAX format. This package also provides novel pathway datasets and advanced querying features for R users through the Pathway Commons webservice allowing users to query, extract, and retrieve data and integrate this data with local BioPAX datasets.

**Availability:** The PaxtoolsR package is compatible with R 3.1.1 on Windows, Mac OS X, and Linux using Bioconductor 3.0 and is available through the Bioconductor R package repository along with source code and a tutorial vignette describing common tasks, such as data visualization and gene set enrichment analysis. Source code and documentation are at http://bioconductor.org/packages/release/bioc/html/paxtoolsr.html. This plugin is free, open-source and licensed under the GNU Lesser General Public License (LGPL) v3.0.

## 1 INTRODUCTION

The amount of biological pathway data in machine-readable databases and formats continues to increase. Pathway analysis allows researchers to gain new understanding of the functions of biological systems. A common task has been to aggregate pathway data across databases. This task has been simplified through the creation of standardized data representations, such as the Biological Pathway Exchange (BioPAX) format (Demir *et al.*, 2010). Pathway Commons is an ongoing effort to aggregate pathway data over a number of databases supporting the BioPAX notation and webservices to access this data (Cerami *et al.*, 2011). The core component that facilitates the development of projects using data in the BioPAX format, such as Pathway Commons, has been Paxtools, a BioPAX application programming interface (API) written in Java (Demir *et al.*, 2013).

While the R programming language is widely used in areas of computational biology, there is a deficiency in the availability of pathway data provided through R packages. A recent review by Kramer et al. describes 12 R packages for working with pathway data (Kramer *et al.*, 2014, 2013). The majority of these packages – including KEGGgraph, PathView, and ReactomePA – utilize and provide data from KEGG and Reactome. A number of the packages are generic parsers for a variety of formats, including the Systems Biology Markup Language (SBML), KEGG Markup Language (KGML), and BioPAX.

Through the PaxtoolsR package, we extend the literature-curated pathway data available to R users, we provide a number of Paxtools API functions, and provide an interface to the Pathway Commons webservice. Through this interface, PaxtoolsR provides native support for the aggregated Pathway Commons database, including data imported from the NCI Pathway Interaction Database (PID), PantherDB, BioCyc, Reactome, PhoshoSitePlus, and HPRD.

## 2 IMPLEMENTATION AND FUNCTIONALITY

PaxtoolsR is implemented using the rJava R package (http://www.rforge.net/rJava/) which allows R code to call Java methods. While R users could use rJava to directly call methods in the Paxtools library, these tend not to follow typical R language conventions, and therefore, PaxtoolsR simplifies the usage of Paxtools in R. PaxtoolsR implements two main sets of features: 1) functions available through the Paxtools console application and 2) functions available provided through the Pathway Commons webservice. Below, we first describe the main data formats used by the PaxtoolsR package and then describe the functions provided by PaxtoolsR. Additionally, the PaxtoolsR provides a vignette to guide users in using the provided functionality, such as the visualization of networks directly in R using existing R graph libaries, such as igraph (Csardi and Nepusz, 2006) and RCytoscape (Shannon *et al.*, 2013), and combining the analysis of gene expression microarrays with pathway data using gene set enrichment analysis (GSEA) (Subramanian *et al.*, 2005).

### 2.1 Data Formats

There are several primary data formats used by the PaxtoolsR package: BioPAX, SIF, and XML; here we describe the role of each of these formats in the PaxtoolsR package.

#### 2.1.1 BioPAX Format

The BioPAX format is an RDF/OWL-based language described previously and used as the main input format for the functions provided via the Paxtools Java library (Demir *et al.*, 2010, 2013). BioPAX representations for databases aggregated by Pathway Commons can be downloaded from the project website (http://www.pathwaycommons.org). The currently aggregated databases include HPRD, HumanCyc, NCI PID, Panther, PhosphoSitePlus, and Reactome.

#### 2.1.2 Simple Interaction Format (SIF)

The SIF format is a tab-delimited, plain-text network edge list that describes how two molecules are related in a binary fashion, and is generated from BioPAX datasets by searching certain graphical patterns (Babur *et al.*, 2014). The SIF format composed of three columns: PARTICIPANT A, INTERACTION TYPE, and PARTICIPANT B. There are 14 possible interaction types, which are described in the package vignette and additional columns may be appended to each line with accompanying metadata. The conversion from BioPAX to SIF is lossy, but remains useful for applications that require binary interactions, which includes many existing network analysis software tools.

#### 2.1.3 Extensible Markup Language (XML)

BioPAX file validation and search results of Pathway Commons results are returned as R XML (http://www.omegahat.org/RSXML/) objects where further data can be extracted using XPath expressions in R.

### 2.2 Convert, Merge, Validate Local BioPAX Files

A number of BioPAX-related functions are available in PaxtoolsR. These functions can both operate on local BioPAX files and those retrieved from Pathway Commons. PaxtoolsR provides a programming interface for the BioPAX format and for the functions provided through the Paxtools console application. These functions allow importing data into R through the SIF format and conversion of BioPAX files into a variety of formats, including the GSEA gene set format. Functions are also provided to extract subnetworks from BioPAX files and the merging of multiple BioPAX files through a previously described method (Demir *et al.*, 2013). Additionally, PaxtoolsR provides methods to summarize the usage of BioPAX classes and validate BioPAX datasets (Rodchenkov *et al.*, 2013).

### 2.3 Query and Traverse Data from Pathway Commons

PaxtoolsR provides a number of functions for interacting with the Pathway Commons webservice. PaxtoolsR allows users to query Pathway Commons data via two functions. The first involves searching for specific molecular species or pathways of interest, using the searchPc() function. The second is the graphPc() function, which allows users to query subnetworks of interest. Figure 1 shows the usage of the graphPc() command to extract a small subnetwork involving the kinases AKT1 and MTOR. This subnetwork is then converted to a binary SIF network and visualized using igraph in R; this showcases how Pathway Commons data can be easily visualized using existing R packages. The traverse() function allows the extraction of specific entries from BioPAX records, such as the phosphorylation site information from proteins described in a BioPAX dataset.

**Fig. 1.**
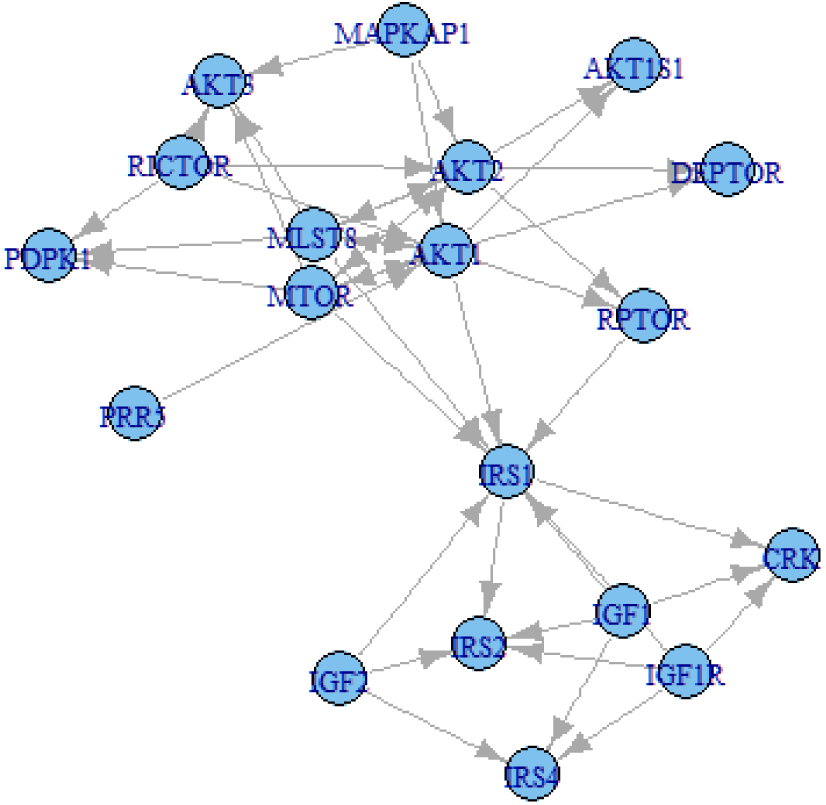
Pathway Commons graph query of interactions between AKT1 and MTOR using PaxtoolsR and visualized using igraph.

## 3 CONCLUSION

The PaxtoolsR package extends the available biological pathway data available to researchers working primarily in an R environment. This package makes many of the features available from the BioPAX Paxtools API and the Pathway Commons webservice. The data and functionality provided here can be used for a wide range of biological pathway analysis studies and can be easily integrated with the rich ecosystem of existing R packages. Future development of this R package is expected as additions are made to the underlying Paxtools Java library and Pathway Commons webservice.

## ACKNOWLEDGEMENT

We would like to thank Alex Root and Eric M. Liu for helpful discussions.

## REFERENCES

Babur, Ö., et al. (2014). Pattern search in BioPAX models. Bioinformatics, 30(1), 139–140.

Cerami, E. G., et al. (2011). Pathway Commons, a web resource for biological pathway data. Nucleic acids research, 39(Database issue), D685–90.

Csardi, G., and Nepusz, T. (2006). The igraph software package for complex network research. InterJournal, Complex Systems, 1695(5), 1–9.

Demir, E., et al. (2010). The BioPAX community standard for pathway data sharing. Nature biotechnology, 28(9), 935–42.

Demir, E., et al. (2013). Using biological pathway data with paxtools. PLoS computational biology, 9(9), e1003194.

Kramer, F., et al. (2013). rBiopaxParser–an R package to parse, modify and visualize BioPAX data. Bioinformatics (Oxford, England), 29(4), 520–2.

Kramer, F., et al. (2014). R-Based Software for the Integration of Pathway Data into Bioinformatic Algorithms. Biology, 3(1), 85–100.

Rodchenkov, I., et al. (2013). The BioPAX Validator. Bioinformatics (Oxford, England), 29(20), 2659–60.

Shannon, P. T., et al. (2013). Rcytoscape: tools for exploratory network analysis. BMC bioinformatics, 14(1), 217.

Subramanian, A., et al. (2005). Gene set enrichment analysis: a knowledge-based approach for interpreting genome-wide expression profiles. Proceedings of the National Academy of Sciences of the United States of America, 102(43), 15545–50.

